# The effect of environmental enrichment on whole-brain gene expression in an imperiled fish

**DOI:** 10.1101/2025.10.18.683224

**Authors:** CJ Kopack, RL Moran, ED Broder, C McDonald, ER Fetherman, KL Hoke, LM Angeloni

## Abstract

When hatchery-reared fish are used to augment wild populations, phenotypic mismatch caused by differences between hatchery and wild environments can limit efforts to conserve fish species at risk of extinction. Some phenotypes adapted to or induced by hatchery environments are thought to be maladapted for life in the wild. Thus, enriching the hatchery environment (abiotically and biotically) to make it more similar to the wild may induce phenotypes, including gene expression profiles, that are better suited to the environments fish will experience after release. Here, we took a molecular approach (TagSeq) to elucidate how abiotic and biotic (predator training) enrichment impacts the whole-brain gene expression of a species of conservation concern, the Arkansas darter (*Etheostoma cragini*), comparing the effects in two hatchery populations to a wild reference population. While we found no effect of biotic enrichment, we found that numerous genes were differentially expressed between populations and abiotic enrichment treatments. Notably, we found that expression profiles of hatchery fish more closely resembled those of wild fish when reared with abiotic enrichment. Functional analysis revealed that many differentially expressed genes were related to feeding behavior, development, and reproduction. These results have implications for conservation, supporting the management of darters at the level of the population and the use of abiotic enrichment to reduce phenotypic mismatch between hatchery and wild fish.

## Introduction

As biodiversity declines, conservation practitioners must creatively use new and existing tools to conserve populations and assess the efficacy of their efforts. One frequently used tool is population augmentation, which involves supplementing populations with individuals from another location or from captivity (Mallinson 1995). However, programs that use captive animals to supplement wild populations often struggle to produce individuals capable of surviving and reproducing after release (Brown and Laland 2001; Brown and Day 2002; Fitzpatrick et al. 2014; Fraser 2008; Griffin et al. 2000; Hawkins et al. 2008; Hutchison et al. 2012; Jackson and Brown 2011; Mallinson 1995; Mesquita and Young 2007; Olla et al. 1998). Stark differences between captive and wild environments may contribute to the poor performance of captive individuals after release into the wild, prompting a critical examination of the effects of the captive environment on phenotypes (Brown and Day 2002; Crane et al. 2015; Evans et al. 2015; Jackson and Brown 2011; Lavoie et al. 2018; Stringwell et al. 2014). This is particularly important for aquatic hatcheries that produce vast numbers of fish for supplementation (Brown and Laland 2001; Brown and Day 2002; Jackson and Brown 2011; Lavoie et al. 2018; Stringwell et al. 2014). The hatchery environment can cause captive fish to differ phenotypically from wild populations through both evolutionary change (Brown and Laland 2001; Brown and Day 2002; Bull et al. 2022; Christie et al. 2012; Christie et al. 2016; Lavoie et al. 2018; Stringwell et al. 2014) and phenotypic plasticity (Brown and Laland 2001; Brown and Day 2002; Crane et al. 2015; Lavoie et al. 2018; Stringwell et al. 2014). The phenotypic mismatch between hatchery and wild fish can be vast (Brown and Laland 2001; Lavoie et al., 2018; Stringwell et al. 2014), encompassing morphology (e.g., Belk 2008; Hutchison et al. 2012; Kihslinger et al. 2006; Saraiva and Pompeu 2016), physiology (e.g., Barcellos et al. 2018; Chittenden et al. 2010; Fuss and Byrne 2002; Hutchison et al. 2012; Salvanes et al. 2013), behavior (e.g., Barcellos et al. 2018; Brown and Day 2002; Crane et al. 2015; D’Anna et al. 2012; Salvanes et al. 2013), life history (e.g., McDermid et al. 2007), and even individual personality (e.g., Johnsson et al. 2014).

One potential tool to address this phenotypic mismatch is to enrich the hatchery environment to better match the wild environments that individuals will experience after release. This may involve enhancing the hatchery environment with abiotic enrichment, including an increase in environmental complexity (D’Anna et al. 2012; Johnsson et al. 2014; Kopack et al. 2023b; Salvanes et al. 2013; Tave et al. 2019; Ullah et al. 2017), and/or biotic enrichment, including experience with foraging and predator avoidance (Brown and Laland 2001; Brown and Day 2002; D’Anna et al. 2012; Kopack et al. 2023a; Kopack et al. 2023b; Sanogo et al. 2011; Vilhunen 2006). Enrichment interventions can cause plastic effects across phenotypes (e.g., morphology, physiology, behavior, etc.), making it challenging to holistically quantify their impacts. Measuring changes in gene expression is one way to reveal plasticity across a broad array of traits in a single assay while also elucidating the mechanisms underlying phenotypic change. Changes in gene expression due to captive rearing environments have been observed in several species, such as cane toads (*Rhinella marina*, Yagound et al. 2022), Nile tilapia (*Oreochromis niloticus*, Konstantinidis et al. 2020), three-spined sticklebacks (*Gasterosteus aculeatus*, Hablutzel et al. 2016), and a variety of hatchery-reared fishes. For example, Salvanes et al. (2013) found that abiotic enrichment increased expression of certain genes in the forebrain related to cognitive performance in hatchery Atlantic salmon (*Salmo salar*). Changes in temperature affected differential gene expression related to metabolism, cell regulation, and signaling between wild and hatchery Australian snapper (*Chrysophrys auratus*; Wellenreuther et al. 2019). Here, we use gene expression to assess the impacts of abiotic and biotic enrichment for a species of conservation concern, the Arkansas darter (*Etheostoma cragini*). While previous studies have used gene expression to identify differences in phenotypes among populations or individuals in different environments, to our knowledge, ours is the first to use gene expression to better understand how we might reduce phenotypic differences between captive animals of conservation concern and their wild counterparts.

The Arkansas darter is a small freshwater fish endemic to North America that is state threatened in Colorado. Current efforts to conserve this species include using hatchery populations to supplement wild populations from which they were originally sourced. However, the genes from stocked fish have not persisted in wild populations, suggesting that supplementation efforts are not effective (Fitzpatrick et al. 2014). We have explored both abiotic and biotic enrichment as possible solutions (Kopack et al. 2023a; Kopack et al. 2023b), with some encouraging results. When combined, the two forms of enrichment caused an increase in survival upon a first encounter with a live predator (Kopack et al. 2023b). However, nothing is known about how abiotic or biotic enrichment affects gene expression in Arkansas darters.

We used Tag-based sequencing (TagSeq; Lohman et al. 2016; Matz 2018) to assess plasticity of gene expression in response to abiotic and biotic enrichment in hatchery-reared darter populations. We also compared gene expression of our hatchery fish to a wild population to determine whether enrichment shifts phenotypes to be more similar to those of wild fish. We addressed three main questions: (1) Does abiotic enrichment affect gene expression of hatchery-reared darters?; (2) Does biotic enrichment (predator training) affect gene expression of hatchery-reared darters?; and (3) If so, do enrichment-induced changes in gene expression make hatchery-reared darters more similar to their wild conspecifics?

## Materials and Methods

### Experimental design

The design of our experiment allowed us to determine whether enrichment affected gene expression of neural tissue in both hatchery-reared and wild Arkansas darters. This design was used in a parallel experiment that examined the effects of abiotic and biotic enrichment on survival in these same darters (Kopack et al. 2023b). Briefly, we reared two populations of captive, hatchery darters in two abiotic treatments (with and without enrichment) to evaluate whether abiotic enrichment affected gene expression. Second, we exposed a subset of these fish to two biotic treatments (with and without predator training) to examine if biotic experience affects gene expression (Figure 1). A wild population not previously exposed to the abiotic treatments was added to the biotic enrichment portion of the experiment to determine if fish in the treatments that mimic the wild environment (abiotic enrichment and predator training) would have gene expression profiles that were more similar to the profiles of wild fish. Following these treatments, we sacrificed darters, collected their neural tissue, and used a Tag-based sequencing approach to identify genes that were differentially expressed among populations and treatments as well as functional analysis.

**Figure 1.**
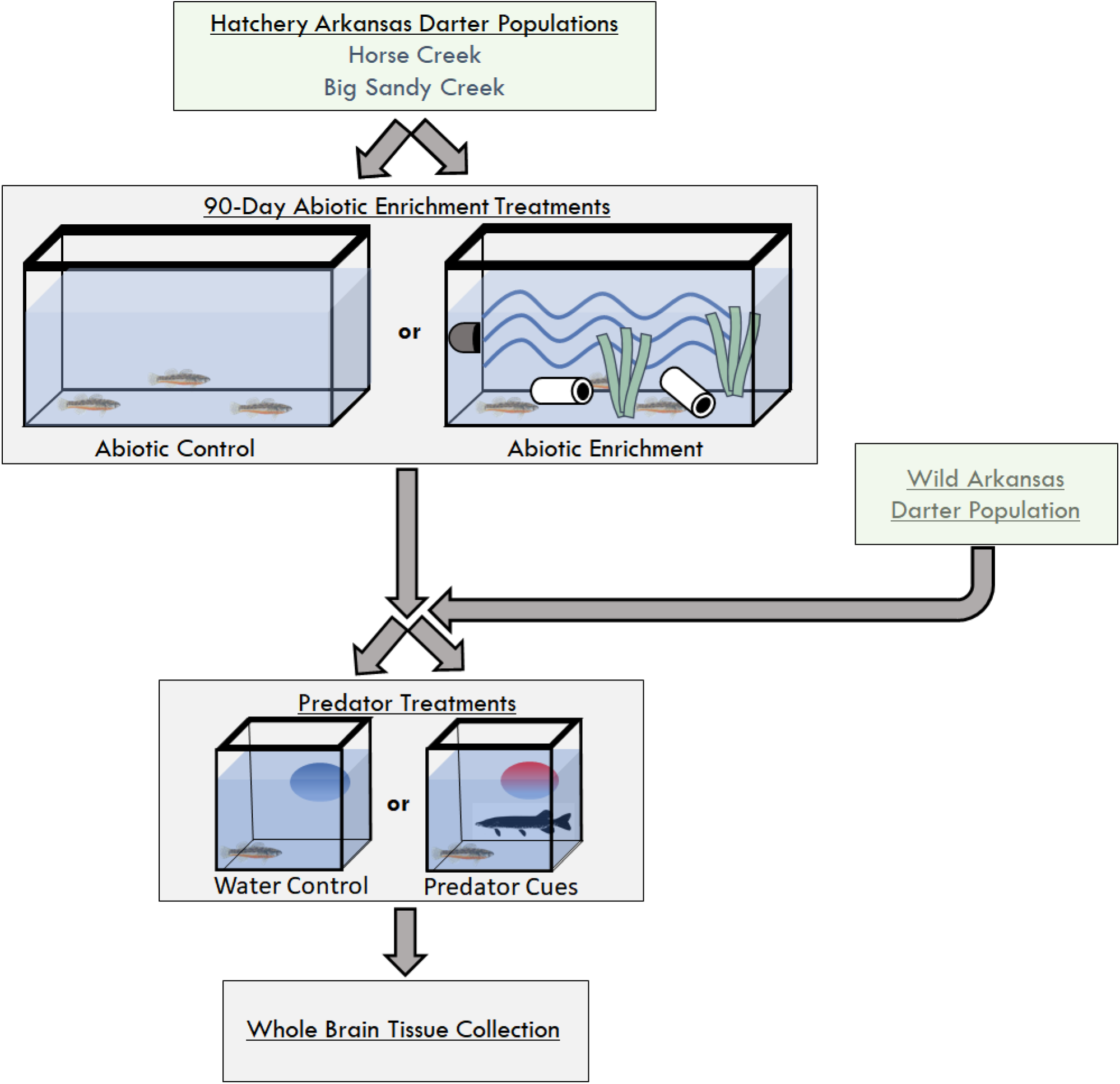
Schematic of experimental design and procedures. Horse Creek and Big Sandy Creek hatchery populations were exposed to abiotic treatments for 90 days. The Abiotic Enrichment treatment tanks included high flows, changing temperatures, artificial plants, and PVC caves. Following abiotic treatments, both hatchery and wild darters were exposed to predator treatments. The Predator Cues treatment included a combination of conspecific alarm cue, predator kairomone, and a predator model. After treatments, whole brain tissues were collected for gene expression analysis.

### Experimental animals

We used three populations of Arkansas darters for this experiment: two hatchery-reared populations, Horse Creek and Big Sandy Creek (named after their original source populations), and a wild population collected from the May Valley Ditch (hereafter ‘Wild’). We obtained 100 young-of-year fish per population that were approximately 10 months old and developmentally mature from the John W. Mumma Native Aquatic Species Restoration Facility (Alamosa, CO) in March 2018, and 20 fish from the May Valley Ditch (Lamar, CO) in June 2018. Hatchery-reared and wild fish were transported to the Colorado Parks and Wildlife (CPW) Salmonid Disease and Sport Fish Research Laboratory (Fort Collins, CO) for experimentation.

### Abiotic and biotic enrichment

The Horse Creek and Big Sandy Creek fish were randomly divided into two abiotic treatments (with and without enrichment) and reared for 90 days (Figure 1; see Kopack et al. 2023b for details). For the control treatment, each population was housed in a separate 76-L flow-through tank supplied with filtered well water at a flow rate of 7.5 L per minute and an average (± SD) temperature of 13.5 ± 2ºC. Abiotic enrichment was added to two additional 76-L tanks, one per population, which included structure (caves made from PVC pipe and artificial plants; multi-pack B1; Marineland®, Blacksburg, VA), temperature variation (daily fluctuations of 3 ± 0.5ºC created by two 100 W in-tank heaters; Eheim™, Buffalo, NY), and flow (two Hydor powerheads 155 L per minute; Hydor USA Inc., Sacramento, CA). The lab was illuminated by 32 W fluorescent lights (General Electric Electorlux) with a light cycle of 14:10 hours light:dark.

Darters were then exposed to biotic enrichment treatments that consisted of either a predator training treatment or a control (Figure 1). We randomly selected darters from the Wild population and from both abiotic treatment groups of the two hatchery populations (Horse Creek and Big Sandy Creek) and randomly assigned them to one of two predator treatments (treatment and control) such that there were five darters per population and treatment combination (Table 1). Twenty-four hours before administering the predator treatment, we moved darters to individual 10-L flow-through tanks receiving the same filtered well water (see Kopack et al. 2023b). Then the control treatment received a water control while the predator treatment received three stimuli simultaneously: 5 mL of conspecific alarm cue, 5 mL of predator kairomone (tiger muskie [*Esox masquinongy* x *E. lucius*]), and a 102 mm 3D tiger muskie model (Savage Gear USA, Ontario, CA; following Kopack et al. 2023a). Alarm cue was extracted from decapitated darters by scoring the skin with a razor blade and rinsing with distilled water (following Kopack et al. 2023a, adapted from Nordell 1998), and predator kairomone was collected from 19-L non-circulating tanks that housed two adult tiger muskie that were fed darters and held for 48 hours (following Kopack et al. 2023a; Kopack et al. 2023b). The 3D model was attached to a wooden dowel via fishing line so that the researcher could place it in the individual darter tank from behind a blind. A researcher administered the predator treatments (water control and predator cues with predator model) from behind a blind and waited 5 minutes before removing the model and turning on the flow-through system to flush any residual cues from the tanks.

**Table 1.** Number of individual Arkansas darters (N=50 attempted, 5 per cell) from which we recovered RNA by population and enrichment treatments. Note that many of the samples had quantities of RNA that were below a threshold to analyze.

|  | Wild | Horse Creek |  | Big Sandy Creek |  |
| --- | --- | --- | --- | --- | --- |
|  |  | Abiotic Enrichment | Abiotic Control | Abiotic Enrichment | Abiotic Control |
| Predator Cues | 5 | 5 | 4 | 0 | 5 |
| Water Control | 4 | 2 | 2 | 0 | 3 |
| Total | 9 | 13 |  | 8 |  |

### Neural tissue collection (whole brain samples)

We collected whole brains from five darters per population and treatment combination (Table 1). We focused on gene expression in brain tissue because this is the organ that is directly involved with changes in behavioral phenotypes (Salvanes et al. 2013). Exactly one hour after biotic enrichment, each individual fish was netted out of its tank, placed on a sterilized (90% ethanol) surface, decapitated, followed by pithing of the brain. Next, we cut the head along its bilateral axis and used sterile forceps (90% ethanol) to collect brain tissue from the cranium, which we immediately transferred into RNAlater (Ambion; Lohman et al. 2016) and stored at - 20ºC until RNA extraction (55 to 62 days later). All tissues were collected within 60 seconds of netting fish out of their treatment tank.

### TagSeq sample preparation and cDNA library synthesis

We used an RNeasy ® Lipid Tissue Mini Kit (cat. No. 74804; Qiagen) to extract RNA from whole brains, following the standard protocols provided with the kit. Quality of RNA isolated was quantified using a Qubit 2.0 fluorometer (Invitrogen; Life technologies). However, we failed to recover RNA from individuals within the abiotic enrichment treatment of the Big Sandy Creek population (many of the samples had quantities of RNA that were below a threshold to analyze); thus, abiotic treatment inferences could only be made for the Horse Creek population (see Table 1). Isolated RNA samples with concentrations of 10 μl or more were then sent to the University of Texas Austin Genome Sequencing and Analysis Facility (GSAF) where they were sequenced on a single lane of an Illumina HiSeq 2500, 2×100 (following Lohan et al. 2016 and Aglyamova et al. 2019).

### Gene expression data preparation and analysis

We assessed sequencing quality using FastQC v. 0.11.9 and multiQC v. 1.12 (Andrews 2010; Ewels et al. 2016). Following quality control, we aligned reads to the published *E. cragini* genome (CSU_Ecrag_1.0; GCF_013103735.1; Reid et al. 2021) using STAR v. 2.7.10a. with default parameters and in quantification mode (Dobin et al. 2013). Gene-level counts generated using quantification mode were then compiled and used for downstream analyses.

All analyses were performed using R statistical software (R Core Team 2020). Prior to analysis, we summed counts across columns to account for technical replicates. Next, the count data were prefiltered by removing all genes with a mean count less than ten (following Love et al. 2014). To compare the impact of abiotic and biotic enrichment on gene expression in whole-brain tissues, we conducted a Principal Coordinate Analysis (PCoA) followed by a formal analysis of differentially expressed genes using DESeq2 (Love et al. 2014).

When assessing the effects of abiotic enrichment on gene expression, we initially included darters from the biotic enrichment portion of the experiment that had received just the water control treatment to eliminate any effects of biotic enrichment. However, after discovering that biotic enrichment had no effect on gene expression (see below), we performed a post-hoc analysis for the effect of abiotic enrichment that included individuals from both the water control and predator cues treatment groups. These are the results we report below. To assess the effect of abiotic enrichment, we compared the levels of differentially expressed genes (DEGs) between the two Horse Creek abiotic treatment groups and the Wild population, with population as a factor nested within abiotic treatment. We considered genes with adjusted (Benjamini-Hochberg) p-values less than 0.05 to be significantly differentially expressed (DEGs).

To assess the effect of biotic enrichment, we compared the number of DEGs between biotic treatment groups (Water Control and Predator Cues) and among populations (Horse Creek, Big Sandy Creek, and Wild), with both included as fixed effects in a mixed model. We report post-hoc pairwise comparisons across treatments and populations.

We used the GO Consortium Gene Ontology Enrichment Analysis tool (http://geneontology.org/) to ask whether any categories of biological processes were overrepresented in our set of DEGs. The zebrafish (*Danio rerio*) reference database was used for this analysis (26,353 genes). Fisher’s exact tests were performed to determine whether the number of genes associated with a given ontology were over- or underrepresented in our sets of DEGs relative to the reference database. For the functional analysis, we first compared the abiotic enrichment treatment to the control treatment across Horse Creek, Big Sandy Creek, and Wild fish. Next, to compare differences among populations while controlling for effects of the environment, we chose to compare the two hatchery populations (Big Sandy Creek and Horse Creek) excluding the Wild population.

## Results

Abiotic enrichment had a significant impact on gene expression. Specifically, compared to their counterparts in the Abiotic Control group, hatchery-reared Horse Creek fish in the Abiotic Treatment group exhibited a gene expression profile that was more similar to the Wild population (Figure 2). From the PCoA, the first axis (MDS1) explained approximately 37% of the variation observed and was attributed to differences among groups (Wild, Abiotic Enrichment, Abiotic Control), while the second axis (MDS2) explained only 9% of the variation captured and was attributed to inter-individual variation (Figure 2). Pairwise comparisons using DESeq2 revealed more DEGs between the Abiotic Control and Wild fish than between the Abiotic Treatment and Wild fish (Figure 3) (Fisher’s exact test: P < 0.0001). In total, we found 4,162 DEGs (out of 27,486 expressed genes) between Wild fish and fish from the Abiotic Control treatments, with 1,620 genes that increased in expression and 2,542 that decreased in expression for the Abiotic Control fish (Figure 3; Table S1). Comparing the Wild fish and fish from the Abiotic Enrichment treatments, a total of 398 DEGs were found, with 140 genes increasing in expression and 258 decreasing in expression for the Abiotic Enrichment fish (Figure 3; Table S2). We also found 1,336 DEGs between fish from the Abiotic Control and Abiotic Enrichment treatments, with 405 increasing in expression and 931 decreasing in expression for the Abiotic Enrichment fish (Figure 3; Table S3). Notably, of the 1,336 DEGs between Abiotic Control and Abiotic Enrichment fish, 85% (1,137) of these genes were also differentially expressed between Abiotic Control and Wild fish. Thus, similar genes were differentially expressed in Abiotic Enrichment and Wild fish compared to control fish, lending further support to the idea that abiotic enrichment induces phenotypes that more closely match wild phenotypes.

**Figure 2:**
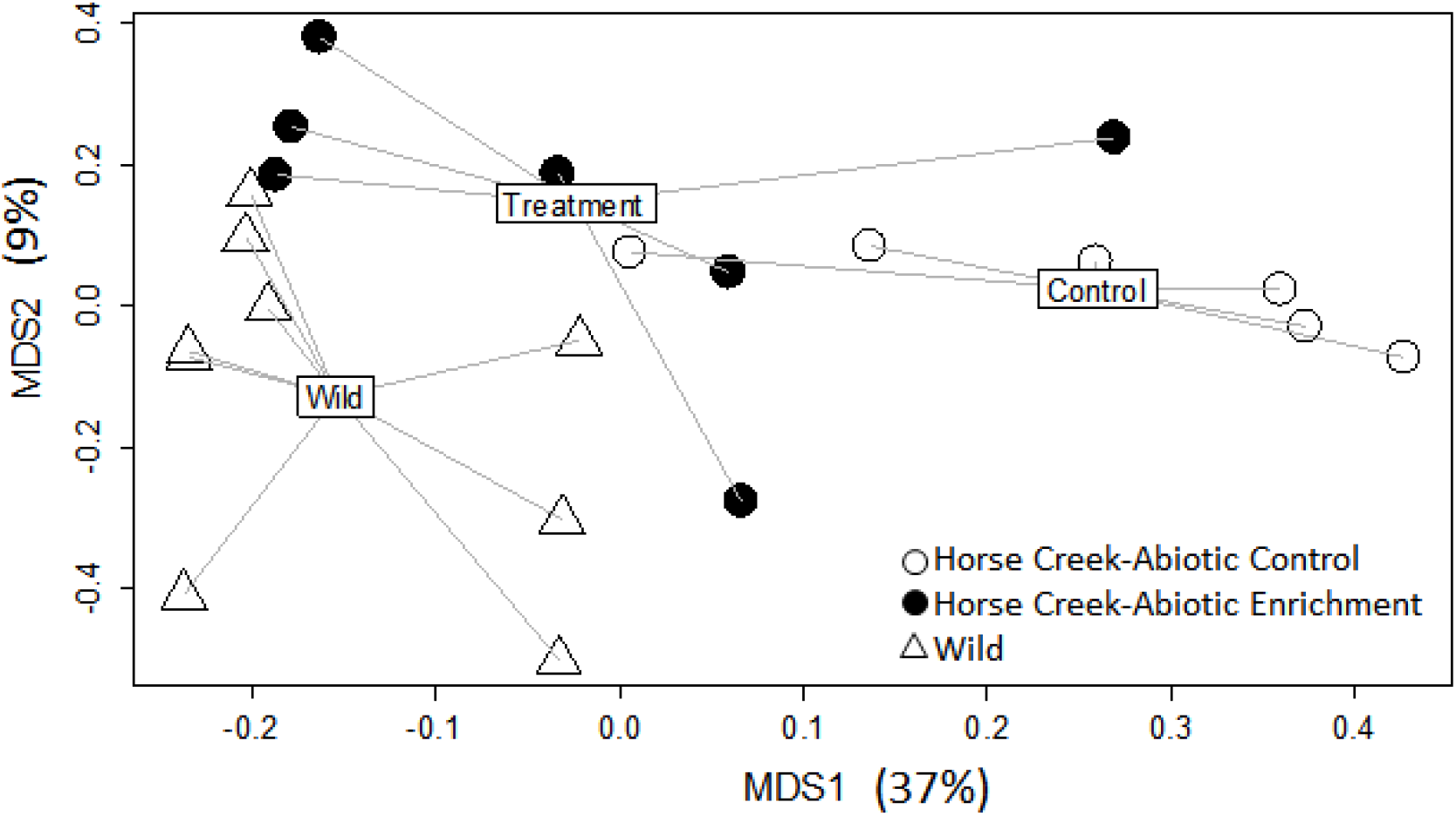
Multidimensional scaling plots showing the gene expression profiles for two populations (triangle = Wild, circle = Hatchery). The Horse Creek hatchery population was exposed to abiotic enrichment treatments (black = Abiotic Enrichment, white = Abiotic Control), whereas the wild population was not (See Figure S1 for a heatmap showing these data).

**Figure 3:**
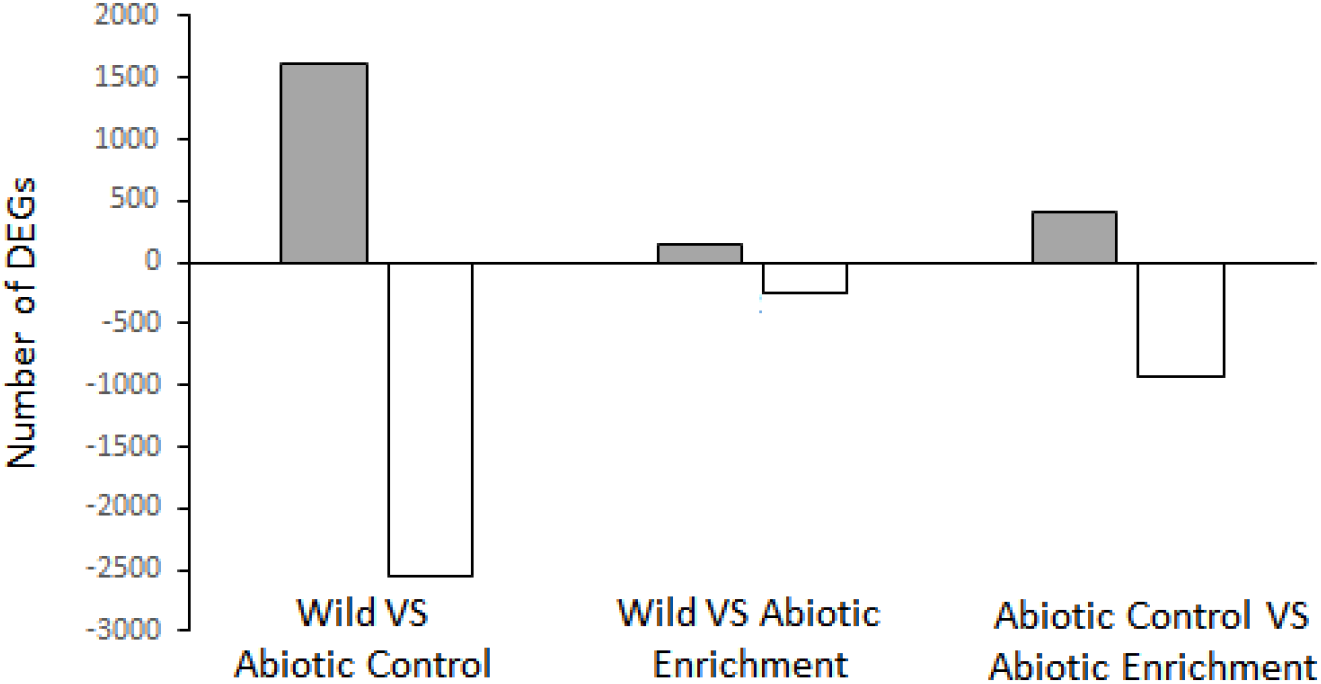
Bar chart showing the number of differentially expressed genes (DEGs) between Wild and abiotic enrichment treatments (Abiotic Enrichment, Abiotic Control). Bars show the number of DEGs that either increased (gray) or decreased (white) in expression for the second experimental group compared to the first.

Genes that were differentially expressed between fish from the Abiotic Control and Abiotic Enrichment treatments were significantly enriched for ontologies related to feeding behavior (*trh*), sensory system development (eye development and morphogenesis: *cdh2, shha, efnb2a, phactr4a*; hair cell differentiation: *cdh2, s1pr2, cops5*), brain structural organization (*cdh2*), metabolism (*atp5f1e, ndufc2, suclg1, suclg1, ndufs2, ndufs7, sdhdb, sdha, cs, sdhb, aclyb, trh, aacs*), hormone regulation (*trh, aacs, ptprna, amh*), and blood vessel development and morphogenesis (*igfbp2a, rack1, nrxn3b, pdcl3, efnb2a, hspa12b, rnf213b, cdh2, eif3i*), just to name a few (Fisher’s exact tests, p < 0.05; Table S8).

The biotic enrichment treatment had no effect on DEGs; however, genes were differentially expressed across populations (Figure 4). From the PCoA, the first axis (MDS1) captured a significant proportion (38%) of the variation and was attributed to differences among populations (Figure 4). The second axis (MDS2) explained approximately 8% of the variation and was attributed to effects of predator training (Water Control vs. Predator Cue treatments; Figure 4), though these effects were not significant. Pairwise comparisons using DESeq2 found zero differentially expressed genes between the fish in the Predator Cues and Water Control treatments (Figure 4; Table S4). However, we did observe a total of 3,093 genes were significantly differentially expressed among the three populations (Figure 5). Comparing the Wild and Horse Creek populations, we found 1,182 of those genes increased in expression while 1,911 decreased in expression for the Horse Creek population (Figure 5; Table S5).

**Figure 4:**
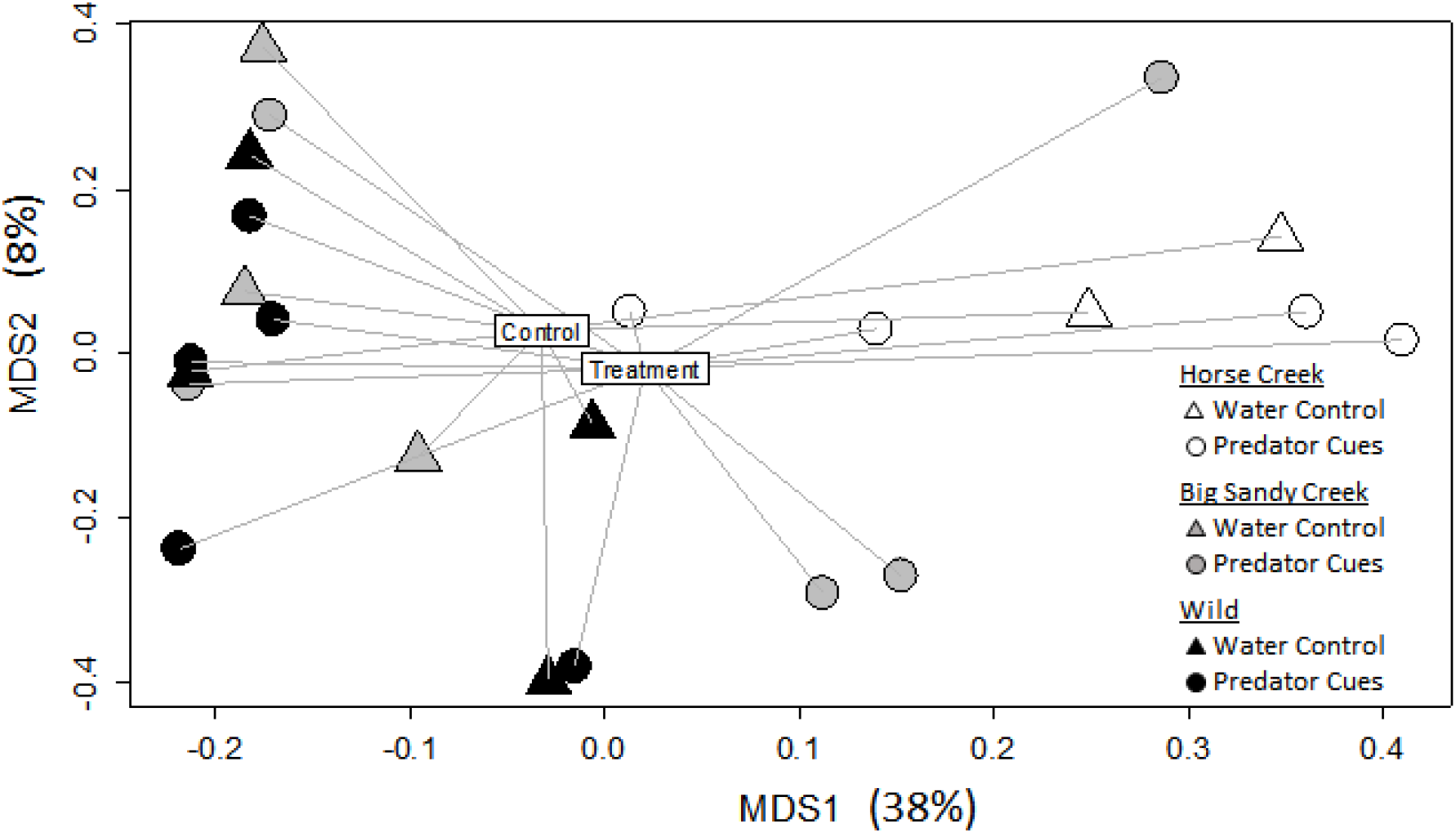
Multidimensional scaling plot showing the gene expression profiles for three populations (black = Wild, gray = Big Sandy Creek, white = Horse Creek) that were exposed to biotic enrichment (predator training) treatments (circle = Predator Cues, triangle = Water Control). See Figure S2 for a heatmap showing these data.

**Figure 5:**
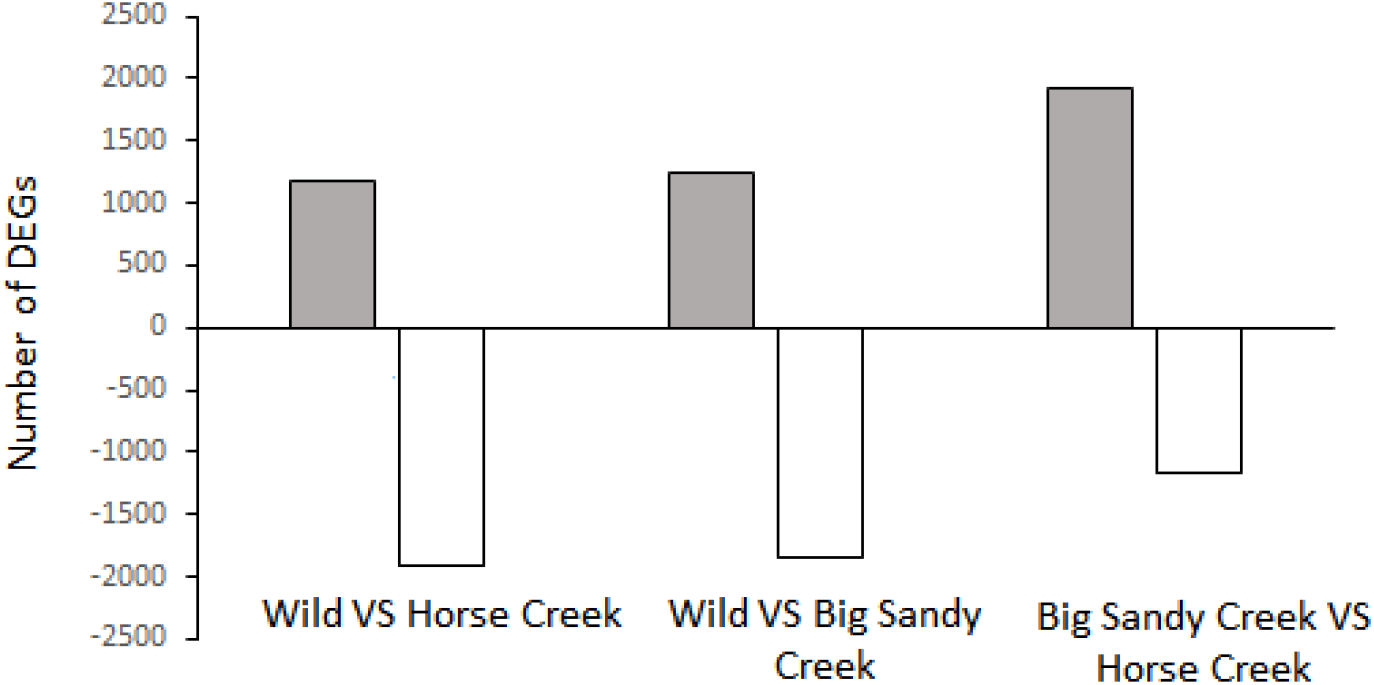
Bar chart showing the number of differentially expressed genes among the three populations (Wild, Horse Creek, Big Sandy Creek). Bars show the number of DEGs that increased (gray) or decreased (white) in expression for the second population compared to the first.

Comparing the Wild and Big Sandy Creek populations, we found 1,246 genes increased expression and 1,847 genes decreased expression for the Big Sandy Creek population (Figure 5; Table S6). Lastly, we found 1,926 genes increased in expression and 1167 genes decreased expression in the Horse Creek hatchery population compared to the Big Sandy Creek hatchery population (Figure 5; Table S7).

Genes that were differentially expressed among populations were significantly enriched for ontologies related to behavior (*oxtr, cdkl5, park7, klhl41a, scn1lab, nop56, trh, atp6v0e1*), sensory systems (*oxtr, oxtr1;* eye development: *lum, vax2, phactr4a, efnb2a*), reproduction (*oxtr, oxtr1*), and embryonic development (*nkx2*.*2a, hspg2, ybx1, notch3*) (Table S9).

## Discussion

In this study, we measured differences in whole-brain gene expression between hatchery and wild populations of Arkansas darters and assessed whether abiotic and biotic enrichment could be used as a conservation tool to reduce those differences. We found that abiotic enrichment caused numerous genes to be differentially expressed between the control and treatment groups. Further, when compared to the wild reference population, the abiotic control group had the greatest number of differentially expressed genes while the abiotic enrichment group had the fewest. In other words, abiotic enrichment induced a plastic response in the Horse Creek population that reflected expression profiles more similar to wild individuals. This suggests that abiotic enrichment may be used to reduce phenotypic mismatch between wild and hatchery fish, potentially priming fish for success in the environment where they will be released. Previous work aligns with this finding. Kopack et al. (2023b) found that abiotic enrichment of hatchery Arkansas darters induced behavioral and morphological responses when implemented alone, as well as increased survival during first encounters with a novel predator when paired with biotic enrichment. Additionally, work in other fish species has shown positive effects of abiotic enrichment for traits of interest, like behavior (Ullah et al. 2017), morphology (Belk 2008), physiology (Chittenden et al. 2010), and even survival in the wild after release (Chittenden et al. 2010; D’Anna et al. 2012).

There are interesting implications concerning the functions of the differentially expressed genes in response to abiotic treatments. First, we detected differentially expressed genes related to feeding behavior, which aligns with prior findings that foraging behavior differed between Arkansas darters in our abiotic treatments (Kopack et al. 2023b). It is also interesting that genes for eye development were differentially expressed. Visual systems have been shaped by sexual selection since visual signals (e.g., male color patterns) are used in female choice, male choice, and male-male competition (Bossu and Near 2015, Hulse et al. 2020, Moran et al. 2017, Moran and Fuller 2018, Williams and Mendelson 2013). Since we observed changes in eye-development genes in our abiotic enrichment treatment (where we changed the environment), it would be interesting to explore the effect of this treatment on female preferences, which would have implications for fitness of both females and males after release in the wild. Importantly, darter visual systems also allow them to navigate their environment, including avoiding predation, so changes in gene expression of the visual system could have played a role in the increased survival during a first encounter with a predator for Arkansas darters that were abiotically and biotically enriched in another study (Kopack et al. 2023b).

We found no evidence that our biotic enrichment induced phenotypic plasticity; there were zero differentially expressed genes between control fish and fish exposed to predator cues. This suggests that the form of predator training we used may not be useful in eliciting observable phenotypic change, supporting previous findings that predator training did not have a strong effect on antipredator behavior in Arkansas darters, perhaps because antipredator behavior is innate (Kopack et al. 2023a). However, this response could also result from the relatively short time fish were exposed to predator cues (reflecting an acute, not chronic, exposure; Snell-Rood 2013) and further research is needed to assess the effects of other forms of predator training (Rittschof and Hughes 2018). For example, Sanogo et al. (2011) induced differences in gene expression by subjecting stickleback fish (*Gasterosteus aculeatus*) to predator cues for longer exposure times during development. Furthermore, the one-hour duration of time between treatment and tissue collection may not have been long enough to detect meaningful changes in expression profiles. However, combined with abiotic enrichment, exposure to predator cues increased survival in Arkansas darters (Kopack et al. 2023b), suggesting there could be an advantage to giving darters experience with predator cues prior to their first encounter with a novel predator. As such, the potential for biotic enrichment to increase survival should continue to be investigated.

We found differences in gene expression among darter populations, with the Horse Creek hatchery population differing most from the Wild population. This suggests that populations differ in regulatory genes with potential impacts on a range of phenotypes and is supported by prior research that found genetic and phenotypic differences among populations (Fitzpatrick et al. 2014; Kopack et al. 2023a; Kopack et al. 2023b). This difference among populations was further supported by our functional analysis, as genes for behavioral and reproductive traits were differentially expressed between hatchery populations. Thus, it may be important to manage darters at the level of the population and maintain genetically separate lineages within the hatchery, as there could be potential adaptive value among them. However, it is unclear if these population differences reflect adaptive differences stemming from the variability between wild or hatchery environments, are a product of drift in their wild source populations, or resulted from bottlenecks or drift in the hatchery setting. Comparisons with the wild populations from which the hatchery populations were sourced could elucidate the origins of these differences. If particular phenotypic profiles are found to be adaptive in the wild, future conservation efforts could choose hatchery individuals with these phenotypes as migrants into wild populations for genetic rescue (Funk et al. 2019) or as targets of selection to enhance captive phenotypes over time (Christie et al. 2016; Dingemanse et al. 2009; Funk et al. 2019).

Future work in the Arkansas darter system should include hatchery-scale enrichment experiments and tests of survival after release. Applying effective abiotic enrichment at the hatchery level should not be logistically difficult; our efforts involved adding simple structure, flow, and temperature variation with readily available materials. It would be useful to explore the duration of abiotic treatment that elicits optimal phenotypic profiles, since darters may spend 10 to 12 months in the hatchery before being released. For instance, we might have elicited an even greater change in gene expression if we had begun the abiotic enrichment right after fish hatched and/or exposed them to the treatment for longer than 90 days. Future research could also investigate the plastic effects of multi-generational enrichment (Christie et al. 2016; Dingemanse et al. 2009; West-Eberhard 1989).

Our finding that abiotic enrichment reduces phenotypic mismatch has implications beyond our study system. Abiotic enrichment in captivity may produce more wild-like phenotypes in other taxa. A few other studies have found differential gene expression in captive animals compared to their wild counterparts (e.g., Nile tilapia, Konstantinidis et al. 2020; cane toads, Yagound et al. 2022; and three-spined sticklebacks, Hablutzel et al. 2016), and abiotic enrichment in captivity has been shown to increase expression of certain genes in the forebrain related to cognitive performance in Atlantic salmon (Salvanes et al. 2013).

We demonstrated the value of measuring gene expression and using enrichment as possible tools in the conservation of an imperiled fish. In doing so, we were able to determine that: 1) abiotic enrichment has the potential to reduce phenotypic mismatch between hatchery and wild populations; 2) darters show no evidence of plasticity in response to biotic enrichment (predator training); 3) there are strong population differences that should be taken into consideration; and 4) there are many candidate genes that potentially underly phenotypic differences among populations, providing valuable information that can be leveraged in conservation efforts. We hope this work inspires others to use gene expression as a tool to measure and reduce phenotypic mismatch between captive and wild animals of conservation concern, and we encourage others to consider novel applications of gene expression to solve conservation problems.

## Acknowledgements

We thank S. Fitzpatrick, Y. Kanno, C. Ghalambor, L. Stein, M. Matz, H. Crockett, P. Foutz, G. Schisler, A. Treble, T. Smith, the Sloan lab, and the staff of the Colorado Parks and Wildlife John W. Mumma Native Aquatic Species Restoration Facility and Wray Fish Hatchery for providing fish, equipment and supplies, logistic support, and expertise. The authors would also like to thank B. Avila, R. Cheek, E. Vigil, T. Swarr, R. Osmundson, T. Hubbard, K. Dick, F. Horne, and M. Whedbee for help with setup, animal husbandry, and data collection.

Funding for this work was provided by Colorado Parks and Wildlife’s Species Conservation Trust Fund (Grant no. SCTF911C) and a Graduate Research Fellowship awarded to CJK by the National Science Foundation. Personnel were supported by NSF grant to EDB (IOS 2240950).

## Ethics approval statement

Fish were cared for under the guidelines and requirements of the Colorado State University Animal Care and Use Committee (Protocol no. 16-6518A).

## Data availability statement

Scripts are archived in a publicly available Zenodo repository (https://doi.org/10.5281/zenodo.7775122). Upon acceptance, sequencing data will be made publicly available via NCBI SRA #XXXXX.

## Benefits Generated

Benefits from this research accrue from the sharing of our data and results on public databases as described above. We also shared our data and results with Colorado Parks and Wildlife as they are the group primarily responsible for the conservation of the Arkansas darter in Colorado.

## Author contributions

CRediT (https://credit.niso.org/): CJK: conceptualization (lead), data curation (lead), formal analyses (lead), funding acquisition (lead), investigation (lead), methodology (lead), project administration (lead), software (lead), visualization (lead), writing – original draft (lead), writing – review and editing (lead); RLM: data curation (equal), formal analyses (equal), methodology (equal), writing – review and editing (supporting); EDB: conceptualization (equal), methodology (equal), writing – original draft (equal), writing – review and editing (equal); CAM: data curation (equal), software (equal), writing – review and editing (supporting); ERF: conceptualization (equal), methodology (equal), resources (equal), writing – original draft (supporting), writing – review and editing (equal); KLH: data curation (supporting), formal analyses (supporting), visualization (supporting), writing – review and editing (supporting); LMA: conceptualization (equal), methodology (equal), writing – original draft (supporting), writing – review and editing (equal).

