## Supplementary figures and images for "The effect of environmental enrichment on whole-brain gene expression in an imperiled fish"

### Supplemental Figure 1

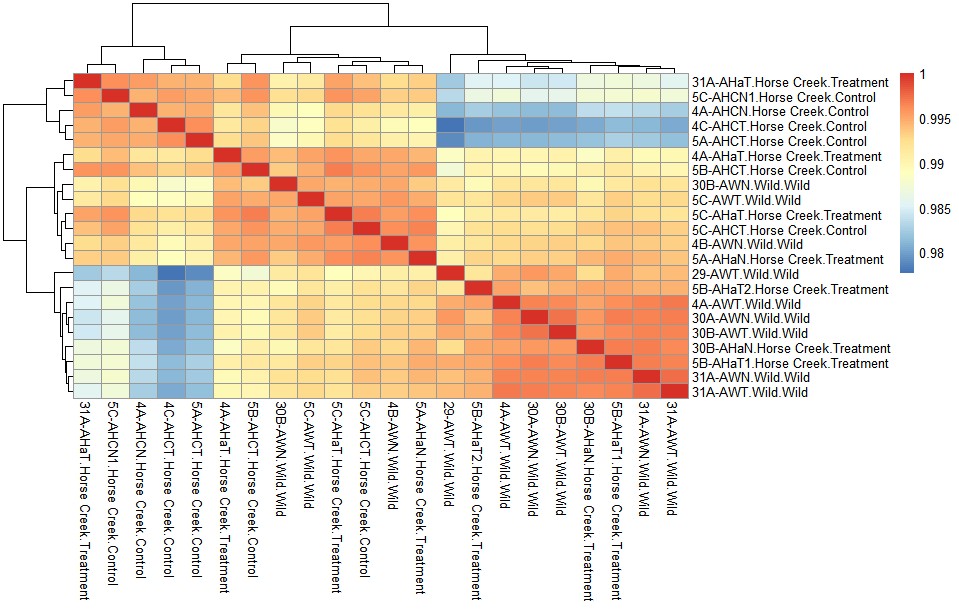

### Supplemental Figure 2

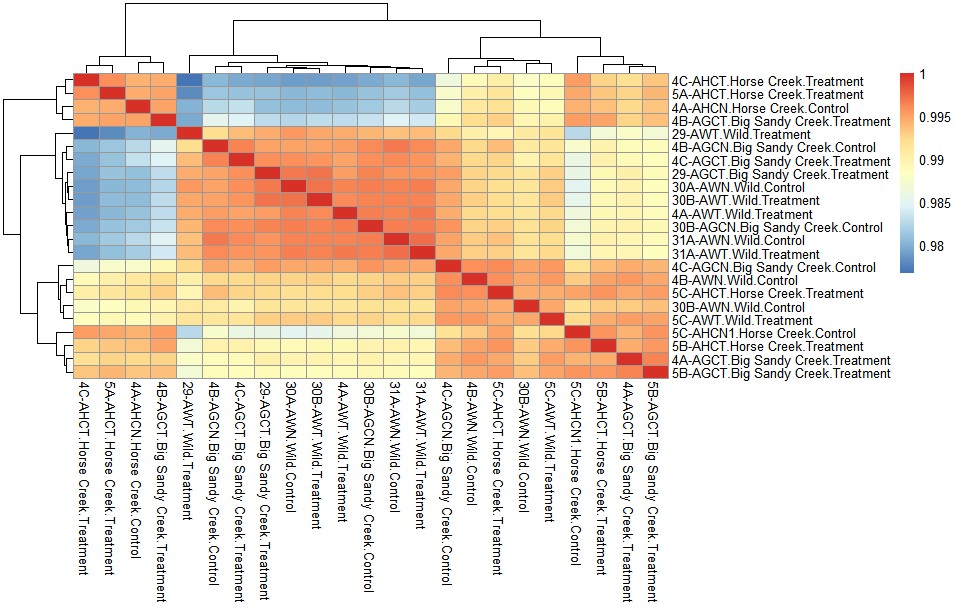
